# Adaptive Representations for Real-World Enzyme Optimization

**DOI:** 10.64898/2026.11.26.768777

**Authors:** Saumya Thakur, David Ming Segura, Philippe Schwaller

## Abstract

Protein fitness landscapes are highly rugged, giving rise to complex optimization landscapes. Bayesian optimization (BO) with Gaussian processes (GPs) is widely used in protein engineering for its sample efficiency and calibrated uncertainty estimates. Here, we study a joint fine-tuning framework of a protein language model (PLM) and a GP across enzyme fitness landscapes that span various distinct mutation regimes encountered in real-world engineering campaigns. Joint fine-tuning optimizes the PLM and GP parameters together in an uncertainty-calibrated manner, providing an adaptive representation that utilizes the information from previously acquired points in addition to the existing knowledge from pretraining. Across seven combinatorial enzyme landscapes, this approach yields a median relative improvement in top-5% coverage of 45% over one-hot encoding and 120% over mean-pooled static ESM-C embeddings. We further explore several new pooling strategies beyond standard mean pooling within the joint fine-tuning framework and show that mutation-site pooling, which localizes the representation to mutated residues, is the most consistent performer across all landscapes, outperforming both established baselines and alternative pooling schemes. Finally, we show that the finetuned approach makes more effective use of training-split information, yielding better coverage across test-split BO rounds. These results suggest that adapting the representation during a campaign, rather than fixing it in advance, is the better design choice for BO-guided enzyme engineering.

## 1 Introduction

The discovery of novel proteins with optimized functional properties is central to modern biotechnology, but remains difficult [Khersonsky and Fleishman, 2022, Listov et al., 2024]. Traditional protein engineering relies on experimental directed evolution [Arnold, 1998], a process that is often prohibitively expensive and time-consuming given the vastness of sequence space and the limited throughput of high-fidelity wet-lab assays. To navigate such landscapes efficiently, Bayesian optimization (BO) [Kushner, 1962, Garnett, 2023] has emerged as a powerful framework, using surrogate models such as Gaussian processes (GPs) [Rasmussen and Williams, 2006] to propose high-fitness candidates in a sample-efficient manner [Yang et al., 2019, Wittmann et al., 2021].

The success of BO, however, depends heavily on the underlying protein representation [Soldát and Kléma, 2024, Ober et al., 2025]. Standard approaches commonly use one-hot encodings [Wu et al., 2019], physicochemical (AAIndex) encodings [Hu et al., 2023], or protein language model (PLM) embeddings [Wittmann et al., 2021, Soldát and Kléma, 2024], the latter benefiting from rapid progress in pre-trained language models for proteins [Rives et al., 2021, Lin et al., 2023]. These representations differ in the prior they encode, but each is fixed before the campaign begins and remains unchanged as fitness measurements accumulate. The limitation is most consequential under mutational shift: campaigns rarely sample a landscape uniformly, instead saturating a region such as the active site during initial screens before extrapolating to distal positions or higher-order combinations of observed hits [Johnston et al., 2024]. A representation fixed in advance encodes no information about which positions the campaign has identified as functionally relevant, leaving the surrogate least informed where extrapolation is most demanded.

In this regard, we make the following contributions:

- *Adaptive representations for real-world enzyme fitness landscapes*. We evaluate enzyme fitness landscapes decomposed by mutation multiplicity (single-vs. multi-mutants), proximity to the active site (close vs. far), and combinatorial complexity, and show that adaptive deep-kernel representations outperform widely used baselines (e.g., one-hot or mean-pooled PLM embeddings) across all of these landscapes.
- *Transferability across distinct mutational landscapes*. Beyond direct discovery, we show on alpha-amylase and TrpB splits that the adaptive representation transfers more reliably under mutational shift than static alternatives.
- *PLM-specific pooling strategies in the adaptive setting*. We explore pooling strategies beyond the standard mean pooling. While joint fine-tuning consistently outperforms established baselines across all of them, mutation-site pooling is the most consistent performer across all landscapes and mutational contexts we test, while a weighted, learned pooling also remains competitive.

## 2 Background and Related Work

### 2.1 Bayesian optimization in biology

Bayesian optimization (BO) is a framework for navigating expensive, high-dimensional search spaces of the kind inherent to tasks with costly evaluations [Kushner, 1962]. BO has become a standard approach in protein engineering, complementing traditional directed- and continuous-evolution methods [Romero et al., 2013, Hu et al., 2023, Liu et al., 2026]. By using a surrogate model, BO provides a principled mechanism for balancing exploration of novel sequences against exploitation of known high-fitness regions. Such surrogate models have traditionally relied on stationary kernels (e.g., Matérn or RBF) paired with simple representations such as one-hot encodings. While effective on simple fitness landscapes, these approaches often struggle in real-world scenarios, since the underlying representation space need not have the geometry required for effective GP optimization. This representational mismatch motivates studying adaptive, PLM-based representations within BO campaigns.

### 2.2 The advent of protein language models

The shift from simple sequence alignments to self-supervised learning has enabled the extraction of rich evolutionary and structural information from massive sequence databases such as UniProt. Protein language models (PLMs) are one such application, adapting the general pre-training recipes used for foundation models to protein sequences, typically via a masked language modeling objective. A range of PLMs have been shown to embed evolutionary knowledge in their latent spaces, most commonly ESM2-650M [Lin et al., 2023], the ESMC family [Hayes et al., 2025], and CARP-640M [Yang et al., 2024]. While these static embeddings are powerful for zero-shot tasks, they are not inherently optimized for specific functional labels such as thermal stability or catalytic activity. Consequently, when used as regression inputs, distances in these latent spaces may not align with the functional gradients of the fitness landscape, creating a disconnect between sequence similarity and functional similarity.

### 2.3 Deep-kernel-based Bayesian optimization

Protein fitness landscapes are characterized by extreme sparsity and high-order epistasis. Traditional surrogate models often struggle with such landscapes because their underlying representations are not optimized for the task’s specific geometry. GPs are one class of surrogate model defined by a kernel function that encodes a notion of distance between inputs (e.g., the *l*_2_ distance between latent vectors). To bridge the gap between discrete sequence space and functional fitness, one can move beyond static distance metrics toward deep kernel methods, in which the geometry of the latent manifold is learned jointly with the surrogate model. This approach extends beyond the limitations of stationary kernels, which do not adapt to better represent the geometry of the fitness landscape.

Most existing methods in this space rely on frozen PLM embeddings or zero-shot priors for a given downstream task [Elnaggar et al., 2021, Meier et al., 2021]. Despite their successes, this limitation remains largely unaddressed. We therefore motivate a joint-optimization regime, in which the representation itself evolves in response to the BO campaign’s objective, and study its effects on a challenging benchmark. This regime creates a bidirectional feedback loop between the PLM and the surrogate, transporting epistemic uncertainty from the GP into the PLM and evolutionary priors from the PLM into the GP. The choice of a GP as the surrogate in this framework is deliberate. Recent work by Chaudhry et al. [2026] demonstrated that MLP ensembles with Thompson sampling can match or exceed GP-based surrogates across a range of protein optimization tasks. Our use of a GP surrogate is motivated by the marginal log-likelihood objective, which jointly calibrates the posterior uncertainty and shapes the learned representation through a single principled loss. While an analogous coupling could in principle be achieved with ensemble-based surrogates, the uncertainty in such models arises from inter-member disagreement rather than from a unified probabilistic formulation, making the integration less straightforward. We leave this extension to future work.

### 2.4 The FLIP2 dataset

To evaluate these methodological advances, the community has moved toward benchmarks that reflect real-world engineering constraints. Released in February 2026, the FLIP2 benchmark [Didi et al., 2026] provides exactly such real-world snapshots of protein engineering campaigns, significantly expanding on its predecessor with datasets spanning enzymes, protein-protein interactions, and light-sensitive proteins. Crucially, FLIP2 implements split strategies (e.g., close-to-far and by-mutation) specifically designed to test a model’s ability to generalize to unseen mutations, a setting in which simpler models can surprisingly outperform conventionally finetuned PLMs. We use this benchmark to quantify the benefits of joint fine-tuning under these realistic, high-extrapolation conditions.

## 3 Methods

In this section, we describe the core methods used in this study, the datasets used for evaluation, and the experiments performed using these methods and datasets. Additional details of the experimental setup and methods can be found in Appendix A.

### 3.1 Gaussian-process-optimized PLMs

GPs are a standard surrogate for sample-efficient optimization, owing to the calibrated uncertainty estimates that provide a principled signal for choosing the next query. A key limitation of GPs, however, is that their knowledge is confined to the experimental data on which they have been fitted. Pre-trained language models, by contrast, carry a rich prior learned from vast pre-training corpora, but are prone to confidently producing incorrect predictions when queried outside that prior. Combining the two approaches, using the GP’s uncertainty quantification to calibrate the language model’s predictions, therefore addresses a weakness of each used alone.

Several recent efforts have built hybrid frameworks that retain the benefits of both components [Ramos et al., 2023, Ranković and Schwaller, 2023]. While most treat the language model and the GP as separate entities, Ranković et al. [2026] and Sin et al. [2026] instead combine the two under a single training objective and show that this joint formulation improves performance over separately trained baselines. In particular, they show that jointly training the language model and GP parameters induces an implicit contrastive-learning effect, yielding a latent space that is more structured with respect to the target property. Here, we extend this idea to enzyme fitness landscapes spanning a range of real-world difficulty levels.

We treat the PLM as a non-linear mapping that projects biological sequences into a latent manifold. Using these embeddings as inputs to a GP surrogate, we distinguish between two regimes: a feature-based approach with static embeddings (equivalent to a stationary kernel), and a representation-learning approach in which a small set of PLM layers is finetuned jointly with the GP parameters. The latter constitutes a deep kernel learning (DKL) framework [Wilson et al., 2016], in which the covariance is defined by composing a base kernel with the transformer (here, a PLM) backbone:

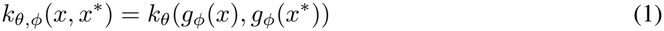

where *k*_*θ*_(·, ·) is a stationary base kernel (*e*.*g*., RBF or Matérn) parameterized by the GP hyperparameters *θ* (e.g., lengthscale, signal variance), *g*_*ϕ*_ : *X* → ℝ^*d*^ is the learned feature map parameterized by the finetuned PLM weights *ϕ, x* and *x*^*\**^ are input biological sequences, and the composition *k*_*θ*_ *g*_*ϕ*_ induces a non-stationary covariance structure over the original sequence space *X*. We adopt a joint-training regime that fine-tunes both the PLM parameters *ϕ* and the GP parameters *θ* by maximizing the marginal log-likelihood. Fine-tuning is performed with Low-Rank Adaptation (LoRA) [Hu et al., 2022], a parameter-efficient technique that substantially reduces the hardware cost of fine-tuning while preserving task performance. Across all experiments, we use Upper Confidence

**Figure 1.**
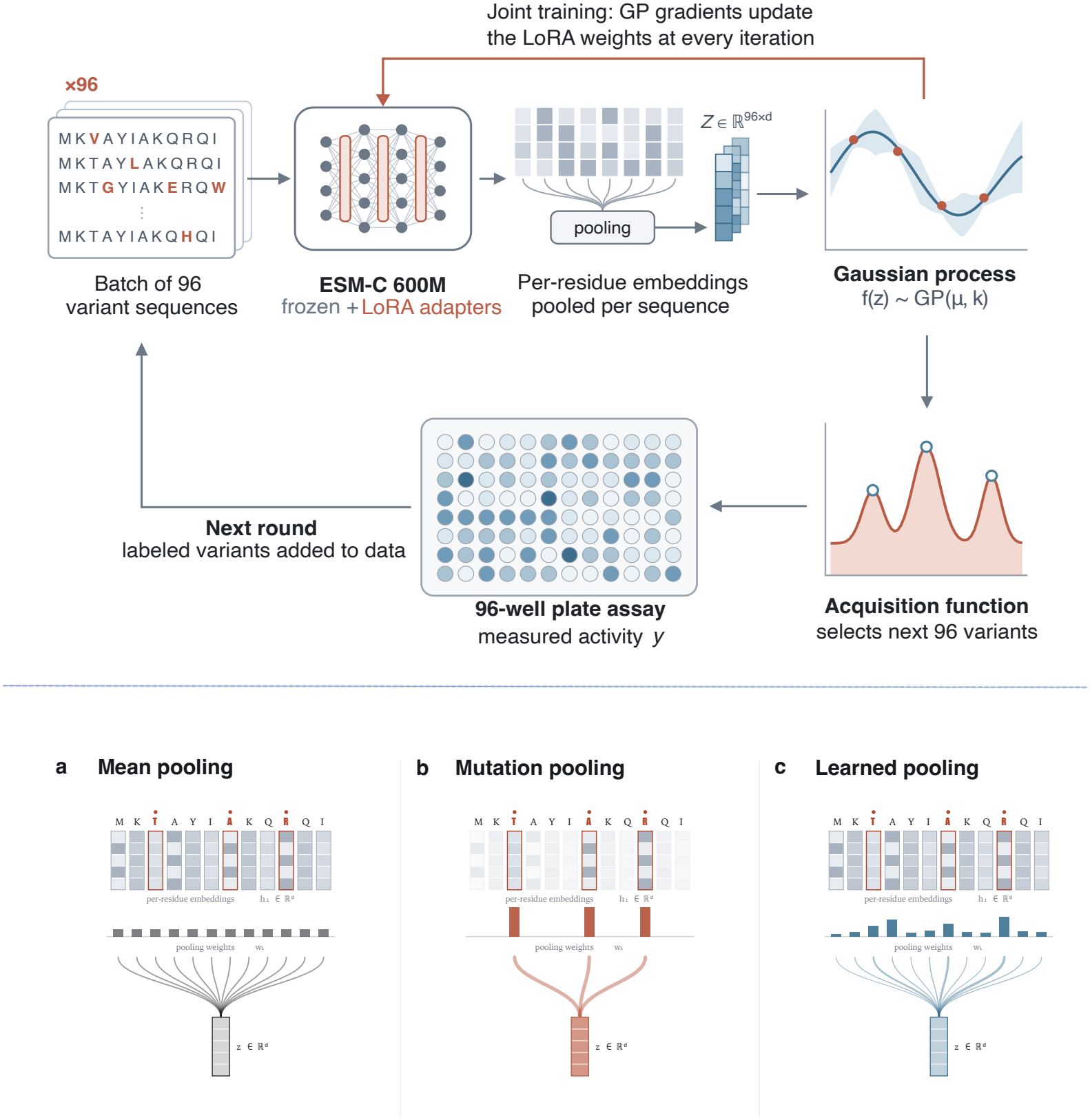
Closed-loop enzyme optimization with a GP-guided LoRA-adapted protein language model. *Top:* each round starts with a batch of 96 variant sequences. The sequences pass through ESM-C 600M, whose base weights stay frozen while small LoRA adapters are trained. Per-residue embeddings are pooled into one vector per sequence, giving a batch embedding matrix. The gradients of the GP marginal log-likelihood flow back into the LoRA weights at every iteration, so the representation and the surrogate are trained together. An acquisition function then picks the next 96 variants, which are measured in a 96-well plate assay. The labelled variants are added to the training data and the loop repeats. *Bottom*: three ways of pooling per-residue embeddings *h*_*i*_ ∈ ℝ^*d*^ into a sequence embedding *z* ∈ ℝ^*d*^. (a) Mean pooling weights every residue equally. (b) Mutation pooling keeps only the mutated positions, shown in red. (c) Learned pooling uses weights *w*_*i*_ trained end to end, so the model decides how much each residue contributes.

Bound (UCB) [Auer et al., 2002] with *β* = 1 as the acquisition function, and initialize the mean and covariance modules as a constant mean and a Matérn-5/2 kernel [Stein, 1999], respectively. The seed set in each campaign is restricted to variants with fitness below the dataset median, which prevents an unrealistically favorable seed set from inflating coverage, and initial samples are drawn via Sobol sampling [Burhenne et al., 2011] to ensure a diverse, well-distributed starting point for GP training.

### 3.2 Dataset

We structure our experiments around seven biologically distinct enzyme fitness landscapes spanning a range of functions, drawn primarily from the FLIP2 benchmark [Didi et al., 2026]. These landscapes span four wild types corresponding to the enzyme families *alpha-amylase, imine reductase, tryptophan synthase β-subunit*, and *aliphatic amidase*. Rather than optimizing over an entire wild type, we restrict each landscape to a subset characterized by a specific mutation depth and position, mirroring the constraints of real-world campaigns. We further apply two split strategies across these landscapes, designed to progressively stress-test generalization under realistic engineering conditions:

- Number-based (one-to-many, two-to-many): train on low-order mutants and test on higher-order combinatorial variants, directly probing a model’s ability to extrapolate across epistatic interactions.
- Position-based (close-to-far, far-to-close): partition mutations spatially relative to the active site, testing generalization to structurally distant regions of the protein.

Together, these landscapes and splits provide a diverse and realistic set of challenges representative of real-world engineering campaigns. Exact sequence counts and split compositions are provided in Appendix B.

### 3.3 Pooling methods

The PLM encoder produces a per-residue embedding for each enzyme sequence; to serve as input to the GP kernel, these residue-level embeddings must be pooled into a single fixed-dimensional vector, making the choice of pooling strategy a design decision within our framework. This choice becomes particularly consequential once the PLM is finetuned: because our datasets derive from deep mutational scanning (DMS), where individual variants differ from one another at only a small number of positions, we explore whether pooling strategies that explicitly localize this signal, such as pooling only the mutated residues, can make more effective use of a finetuned representation than the standard practice of mean pooling across the whole sequence. We therefore evaluate a spectrum of pooling methods, with mean pooling and mutation pooling at the two extremes and a learned, weighted pooling in between. Writing **h**_1_, …, **h**_*L*_ ∈ℝ^*d*^ for the per-residue embeddings of a sequence of length *L* and ℳ for the set of mutated positions, the three schemes pool to a single vector **h** as

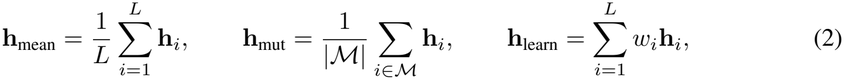

where **w** = softmax(**v**) and **v** ∈ℝ^*L*^ is trained jointly with the PLM and GP parameters. Mean and mutation pooling are thus the two extremes of a fixed-weight family, and learned pooling relaxes the weights to be fit from the fitness signal.

### 3.4 Experiments

We benchmark jointly trained GP and embedding representations across the landscapes described above, using random search as a baseline. Our headline metric throughout is *top-5% coverage*: the fraction of the landscape’s true top-5% variants, ranked by measured fitness, that are contained in the set of variants acquired over the campaign, reported after 5 BO rounds. For the TrpB four-site combinatorial landscape, whose candidate space is roughly an order of magnitude larger than the others, we report top-1% coverage instead. Our experiments fall into two broad setups:

- **Discovery experiments**: We run BO for 5 iterations on each landscape individually and compare representations by top-5% coverage.
- **Generalization experiments**: We evaluate the generalization capability of each representation by scoring the trained GP on the held-out test set and reporting Spearman correlation and recovery of true top-10 (Rec@10). We additionally run BO rounds over the test set to measure downstream discovery performance directly, reporting top-5% coverage on the test set as well.

All experiments use a batch size of 96, consistent with standard 96-well assay sizes in protein engineering campaigns [Auld et al., 2020, Goddard and Reymond, 2004], and are repeated across 10 random seeds to characterize statistical variation.

## 4 Results and Discussion

### 4.1 Discovery performance across splits

LoRA fine-tuning with mutation-site pooling outperforms one-hot and static mean-pooled ESM-C by a median of 45% and 120% respectively, computed as the median over the seven landscapes of the per-landscape relative improvement in coverage (top-1% for TrpB four-site). The mean- and learned-pooled variants also clear both baselines by a wide margin, though mutation-site pooling is the most consistent. Per-landscape coverage is shown in Figure 2.

**Figure 2.**
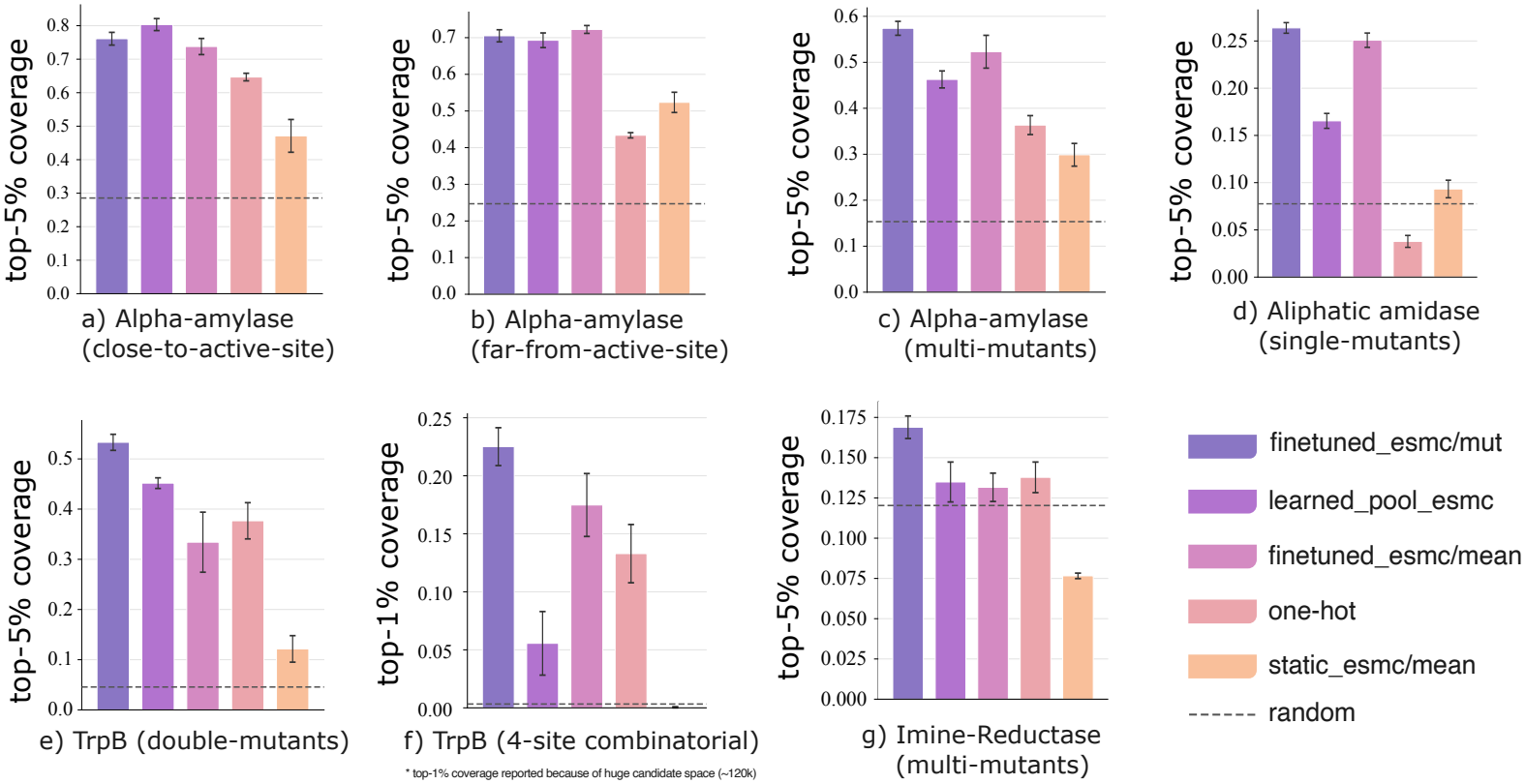
Top-5% coverage after 5 rounds of BO (batch size of 96) across seven enzyme fitness landscapes, comparing LoRA-finetuned ESM-C under mutation-site, learned, and mean pooling against one-hot and static mean-pooled ESM-C baselines, relative to random search (dashed line). Bars show means over 10 seeds; error bars denote SEM. Because the TrpB four-site combinatorial landscape (panel f) has a substantially larger candidate space (~ 120k variants), we report top-1% coverage for this panel instead of top-5%.

In particular, one-hot encoding (OHE) performs worse on the landscape of mutations distal to the active site than on the landscape of mutations close to it (Figure 2), whereas the mutation-pooled and learned-pooled configurations remain consistent across both. One-hot encoding collapses entirely on the aliphatic amidase landscape, which comprises single mutants distributed across the full sequence length. More broadly, static mean-pooled embeddings, despite being widely used in practice for this purpose [Soldát and Kléma, 2024, Meehl and Siddavatam, 2025], trail behind on nearly every landscape we consider.

Although one-hot encoding has been shown to be comparable to PLM-based methods on fitness-prediction tasks across splits [Didi et al., 2026], our results show that in discovery campaigns, where training data is extremely limited, finetuned PLMs provide a substantial advantage over one-hot encoding.

### 4.2 Generalization experiments

The generalization experiments use the position-based and number-based splits of the alpha-amylase and tryptophan synthase datasets to evaluate the surrogate’s fit on the test splits. We report both the Spearman correlation and Rec@10 (recovery of true top-10 candidates in the test set) values as global rank correlation is often a weak proxy for BO performance. The adaptive representation yields the most consistent transfer, as measured by the performance metrics in Table 1. In particular, one-hot and frozen embeddings recover none of the top-10 candidates on either split under position extrapolation, whereas the finetuned representation recovers 18% and 24% respectively.

**Table 1:** Generalization across alpha-amylase and TrpB splits. The GP is trained on candidates acquired from the train split and evaluated for generalization capability on the test set. Mean± SEM over 10 seeds; best per split in bold.

| Split (train → test) | Representation | Spearman *ρ* (↑) | Rec@10 (↑) | Cov (top5/top1 †) (↑) |
|---|---|---|---|---|
| AA (Close → Far) | one-hot | 0.293 ± 0.001 | 0.00 ± 0.00 | 0.113 ± 0.006 |
|  | frozen ESM-C | 0.357 ± 0.009 | 0.00 ± 0.00 | 0.117 ± 0.006 |
|  | finetuned ESM-C (ours) | **0.413 ± 0.019** | **0.18 ± 0.06** | **0.123 ± 0.017** |
| AA (Far → Close) | one-hot | 0.397 ± 0.002 | 0.00 ± 0.00 | 0.144 ± 0.009 |
|  | frozen ESM-C | **0.423 ± 0.021** | 0.00 ± 0.00 | 0.161 ± 0.011 |
|  | finetuned ESM-C (ours) | 0.417 ± 0.006 | **0.24 ± 0.03** | **0.292 ± 0.013** |
| AA (One → Many) | one-hot | 0.467 ± 0.001 | 0.10 ± 0.00 | 0.119 ± 0.002 |
|  | frozen ESM-C | 0.511 ± 0.001 | 0.10 ± 0.00 | 0.117 ± 0.002 |
|  | finetuned ESM-C (ours) | **0.555 ± 0.007** | **0.14 ± 0.02** | **0.276 ± 0.002** |
| TrpB † (Two → Many) | one-hot | 0.220 ± 0.026 | 0.24 ± 0.03 | 0.386 ± 0.036 |
|  | frozen ESM-C | 0.273 ± 0.027 | 0.08 ± 0.03 | 0.147 ± 0.022 |
|  | finetuned ESM-C (ours) | **0.299 ± 0.031** | **0.37 ± 0.04** | **0.446 ± 0.042** |

To further establish the downstream generalization capabilities of the adaptive representation, the top-5% (top-1% for TrpB) coverages after 5 iterations (96 batch size) are also presented in Table 1. Finetuned ESM-C provides the maximum coverage, sometimes even two-fold, when GP is trained only on the train split. Notably, a dissociation between Spearman correlation and coverage is visible on the Far →Close split, where frozen ESM-C attains the best Spearman correlation (0.423) yet barely half the top-5% coverage of the finetuned arm (0.161 vs. 0.292). Running BO directly on the test split confirms the overall pattern: active learning with the finetuned representation transfers better under the mutational shifts typical of real-world campaigns (Figure 3). This indicates that adaptive representations make better use of information acquired under a different mutational distribution in earlier rounds (reflected in coverage at iteration 6), and that finetuned arms transfer more effectively than their cold-start counterparts. Further details are given in Appendix C.

**Figure 3.**
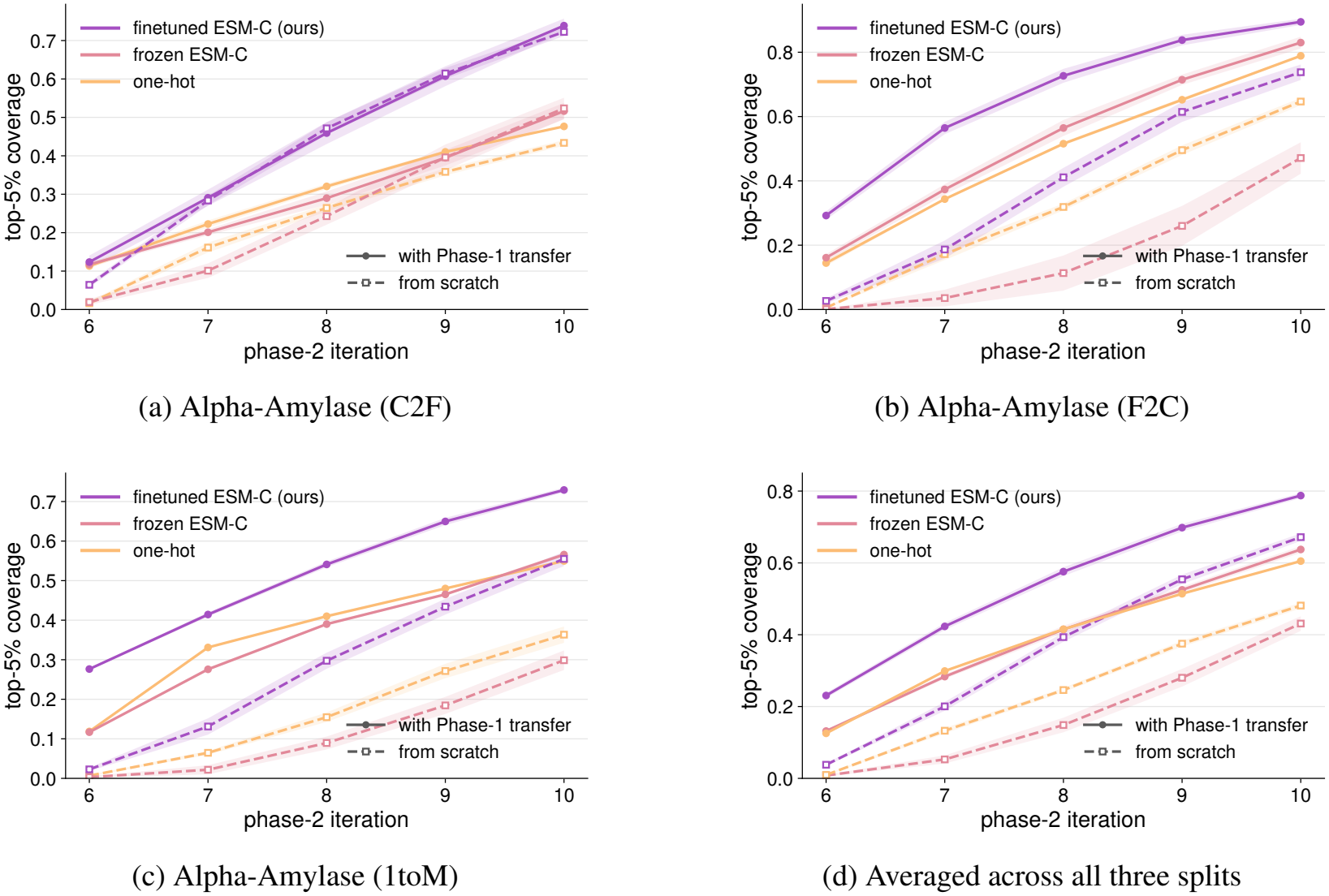
Per-landscape performance difference between cold-start and warm-start BO on the test set (a–c), together with the trend averaged over all three splits (d). Warm-start uses data from the initial rounds of GP fitting; the plots show how different representations make use of data from a different mutational distribution.

## 5 Conclusion

We have explored a joint PLM-GP fine-tuning framework for enzyme discovery campaigns and shown that it consistently outperforms established and commonly utilized representation baselines across a diverse set of real-world mutational landscapes. The advantage holds across mutation depth, proximity to the active site, and combinatorial complexity. One-hot encoding degrades on mutations distal to the active site and collapses on the aliphatic amidase single-mutant landscape, while the adaptive representations remain stable across both regimes. Moreover, mean-pooled static PLM embeddings are consistently found to be underperforming.

The advantage extends beyond in-distribution discovery: the adaptive representation transfers more reliably under mutational shift, continuing to surface top-ranked candidates as the candidate pool moves away from what has already been screened. This provides a practical perspective to experimentalists who are using Bayesian Optimization in their directed evolution campaigns. Additionally, we also evaluate various pooling strategies and demonstrate that mutational pooling is the safest default once the representation is being adapted.

### Limitations

The clearest cost of the approach is the per-round training that adaptation incurs and static baselines do not. Parameter-efficient fine-tuning keeps this within modest hardware budgets, but does not remove it entirely. The trade is favorable where measurements are slow and expensive, as in the low-throughput campaigns targeted here, and less so in high-throughput settings where a static representation evaluated over more rounds may allocate resources better.

Overall, these findings position joint fine-tuning as a practical component of machine-learning-guided enzyme engineering, particularly in the low-data, high-cost regimes where sample efficiency is decisive.

## Acknowledgements

We thank Joshua Sin for constructive discussions and feedback. S.T., D.M.S. and P.S. acknowledge support from the Swiss National Science Foundation Starting Grant (grant number TMSGI2_226509).

P.S. acknowledges support from the NCCR Catalysis (grant number 225147), a National Centre of Competence in Research funded by the Swiss National Science Foundation.

## A Method

The method used in this study is a deep-kernel, Gaussian-process-based Bayesian optimization framework.

### A.1 Bayesian optimization

Bayesian optimization (BO) is a strategy for the global optimization of black-box functions that are expensive to evaluate. We consider the problem of finding a global maximizer:

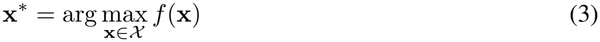

where *X* is the search space (e.g., the space of enzyme sequences in the current study). BO consists of two primary components: a probabilistic surrogate model and an acquisition function.

#### A.1.1 Surrogate model

A surrogate model is a probabilistic approximation of an expensive, complex, or otherwise unknown black-box function. In the context of protein engineering, it acts as a computationally cheap stand-in for the real-world fitness landscape: rather than running a costly wet-lab experiment for every candidate sequence, a BO loop queries the surrogate to predict which sequences are most likely to yield high fitness or high information gain.

Common choices of surrogate model include random forests, Bayesian neural networks, and Gaussian processes. We adopt a Gaussian process (GP), owing to its ability to provide calibrated uncertainty estimates by defining a distribution over functions *f* (**x**) ~ *GP* (*m*(**x**), *κ*(**x, x**^*′*^)), where *κ*(**x, x**^*′*^) is the kernel value between inputs **x** and **x**^*′*^ defined by a kernel function and *m*(**x**) is the mean function. Given a set of *n* observations 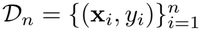, the joint distribution of the observed values **y** and a function value at a new point **x**_*\**_ is:

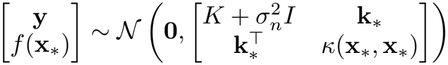

where *K* is the covariance matrix.

The resulting posterior predictive distribution is Gaussian,

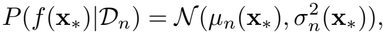

with mean and variance given by:

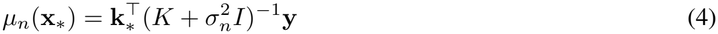

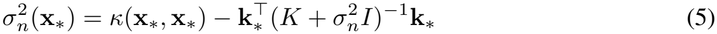

#### A.1.2 Acquisition function

Acquisition functions guide calibrated sequential decision-making within a BO loop. Given the GP posterior, an acquisition score is computed for every candidate sequence, producing a ranking over the candidate space; the top-ranked sequences are then selected for the next round of expensive function evaluations. There are several common choices of acquisition function. Probability of Improvement (PI) [Kushner, 1964] scores a candidate by the chance that it beats the best value observed so far. Expected Improvement (EI) [Jones et al., 1998] scores it by the expected amount it beats that value. Thompson sampling [Thompson, 1933] draws a sample from the GP posterior and picks the candidate that maximizes it. Upper Confidence Bound (UCB) [Auer et al., 2002, Srinivas et al., 2010] adds a multiple of the posterior standard deviation to the posterior mean.

In this work we use UCB, defined as

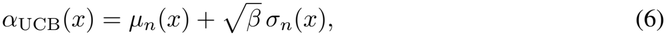

where *µ*_*n*_(*x*) and *σ*_*n*_(*x*) are the posterior mean and standard deviation from Eqs. (4) and (5), and *β* ≥0 sets the trade-off between exploitation and exploration. A small *β* favors candidates with a high predicted fitness. A large *β* favors candidates where the model is uncertain. We set *β* = 1 in all experiments. At each round, we score every candidate in the pool and select the top 96 by *α*_UCB_ as the next batch.

### A.2 Gaussian processes and choice of kernel

For protein engineering, the Matérn-5/2 kernel is the most common choice, as it provides a balance between smoothness and flexibility:

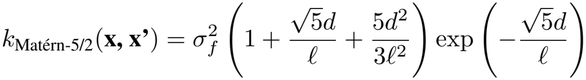

where *d* = ‖*g*_*ϕ*_(*x*) − *g*_*ϕ*_(*x*^*′*^) ‖_2_ is the Euclidean distance in the latent space parameterized by the transformer *ϕ*, and *l* is the learned lengthscale.

### A.3 Implicit contrastive learning via the marginal log-likelihood

Contrastive learning is a paradigm for learning latent representations in which “similar” data points are pulled together and “dissimilar” points are pushed apart. For proteins, we define similarity with respect to the fitness values being optimized rather than evolutionary similarity among sequences. Accordingly, unlike the contrastive objectives used in standard language model pre-training, we use a **fitness-aware contrastive learning objective**, obtained by optimizing the negative marginal log-likelihood (MLL) with respect to the PLM parameters *ϕ*:

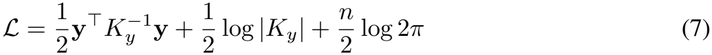

The gradient 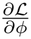 acts as an implicit contrastive force: the quadratic term 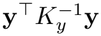 penalizes large latent distances between sequences with similar fitness labels, warping the latent manifold so that out-of-distribution test points are pulled back within the reach of the kernel lengthscale.

Figure 4 shows this warping in action: over successive iterations, high-fitness sequences (light markers) move closer together, as do low-fitness sequences (dark markers), while respecting their target values.

**Figure 4.**
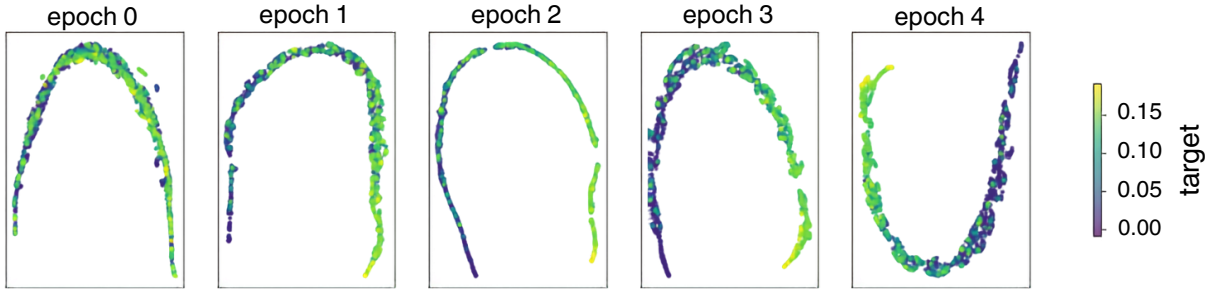
Evolution of the latent space under the implicit contrastive effect obtained using UMAP.

### A.4 Implementation details

#### Static/frozen PLM embeddings

One baseline setting uses static embeddings obtained from a pre-trained language model without fine-tuning. Such embeddings have been shown to encode evolutionary information about protein sequences and are widely used in downstream tasks such as variant-effect prediction. We report results under several pooling strategies applied to these embeddings: mean pooling, mutation pooling (with and without neighborhood context), and learned pooling, described further in Section A.5.

#### PEFT-adapted joint fine-tuning

Parameter-efficient fine-tuning (PEFT) methods adapt large pretrained models to a specific task while updating only a fraction of their weights. By freezing most of the pre-trained network and training only a small set of adapter weights or additional layers, PEFT achieves performance comparable to full fine-tuning at substantially lower hardware cost. We use Low-Rank Adaptation (LoRA) [Hu et al., 2022] to fine-tune the language model, training only the top 25% of its layers. For ESM-C, this corresponds to training 2,067,904 of 577,104,896 total parameters (0.36%), which meaningfully reduces both hardware requirements and per-iteration runtime while retaining the benefits of fine-tuning.

### A.5 Pooling methods

Pooling compresses per-residue attention outputs into a single vector representing the whole sequence.

#### Mean pooling

The most common pooling method, adapted directly from natural language tasks: all token (residue) embeddings are averaged across the sequence to produce a single *d*-dimensional vector representation of the protein.

#### Mutation pooling

Mutation pooling averages only over the mutated positions, motivated by the need to localize the fitness-relevant signal within combinatorial datasets. Because variants in these datasets typically differ from one another at only a small number of positions, mean pooling over the full sequence can dilute the information needed for a meaningful representation.

#### Learned pooling

Learned pooling lies between the two schemes above: we learn a weighted pooling over the full sequence length, with weights fit jointly on the GP training data so that the pooling itself becomes informed by the fitness signal being optimized.

## B Dataset Details

To evaluate the generalization capabilities of the deep-kernel method, we use enzyme datasets from the FLIP2 benchmark [Didi et al., 2026]. These landscapes are specifically designed to test model robustness under the distribution shifts encountered in real-world protein engineering. Note that these shifts are not shifts in the absolute representation space, but rather reflect real-world shifts in the type of mutations a researcher is interested in exploring.

### B.1 Dataset descriptions and splits

We focus our evaluation on four enzyme systems: alpha-amylase (Amylase), tryptophan synthase *β*-subunit (TrpB), imine reductase (IRED), and aliphatic amidase (AMIE). Sequence counts for each split are given in Table 2, and the corresponding fitness distributions are shown in Figure 5.

**Table 2:** Characterization of various datasets used.

| Protein | Subset | #Sequences | Length | Fitness range |
|---|---|---|---|---|
| α-amylase | Single mutants | 488 | 425 | [−0.002, 0.212] |
| α-amylase | Multi-mutants | 3,218 | 425 | [−0.010, 0.201] |
| α-amylase | Close (near active site) | 1,782 | 425 | [−0.010, 0.212] |
| α-amylase | Far (distal to active site) | 1,924 | 425 | [−0.009, 0.191] |
| Aliphatic Amidase | Single mutants | 6,289 | 341 | [−10.52, 0.407] |
| Imine Reductase | Multi-mutants | 4,178 | 290 | [−4.84, 3.01] |
| Tryptophan synthase β subunit | Double mutants | 10,791 | 389 | [−0.676, 1.84] |
| Tryptophan synthase β subunit | Four-site combinatorial | 129,518 | 389 | [−0.402, 2.45] |

**Figure 5.**
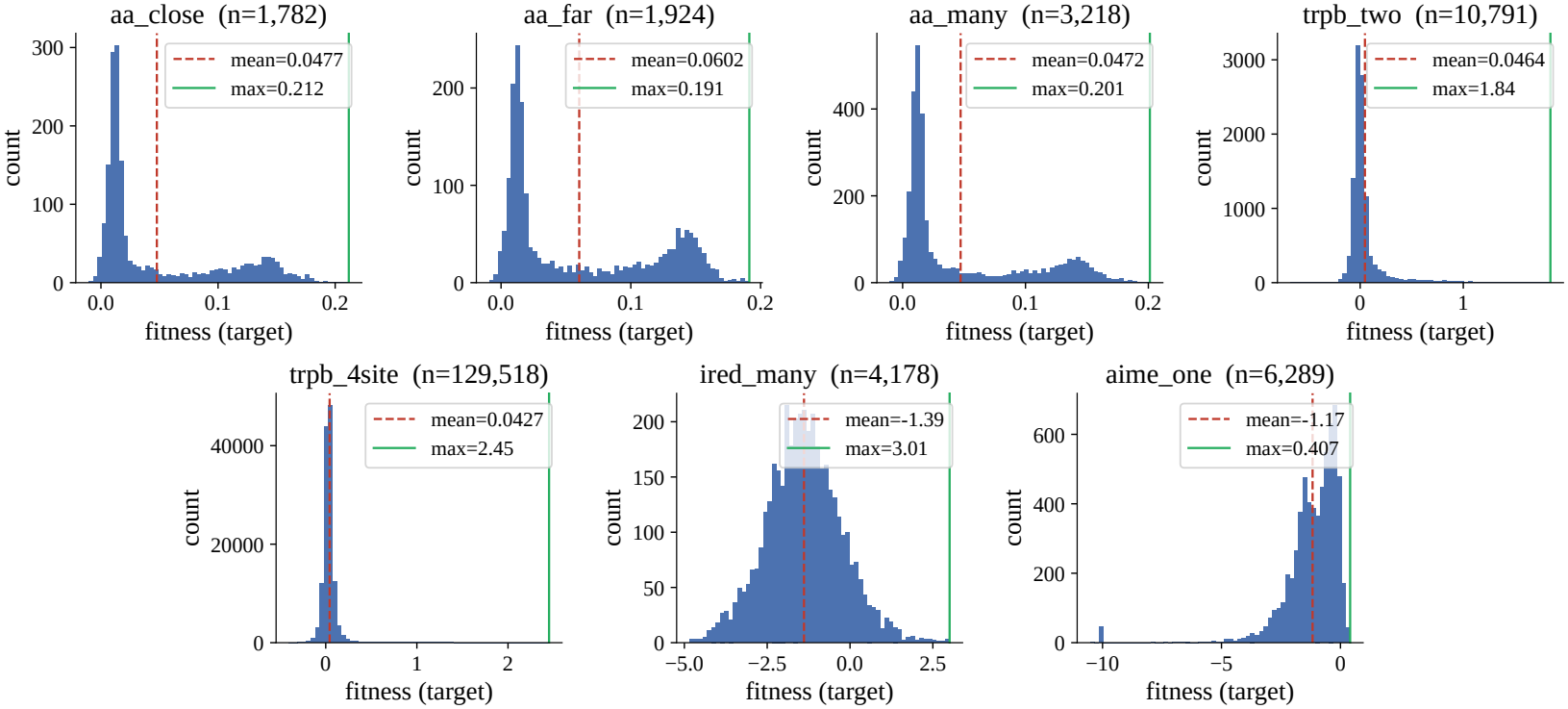
Fitness distribution for all enzyme landscapes

Figure 6 and Tables 4 and 5 label each landscape with the short internal code used in our experiment logs, rather than the descriptive names used in Figure 2; Table 3 maps these codes to their corresponding landscape.

**Table 3:**
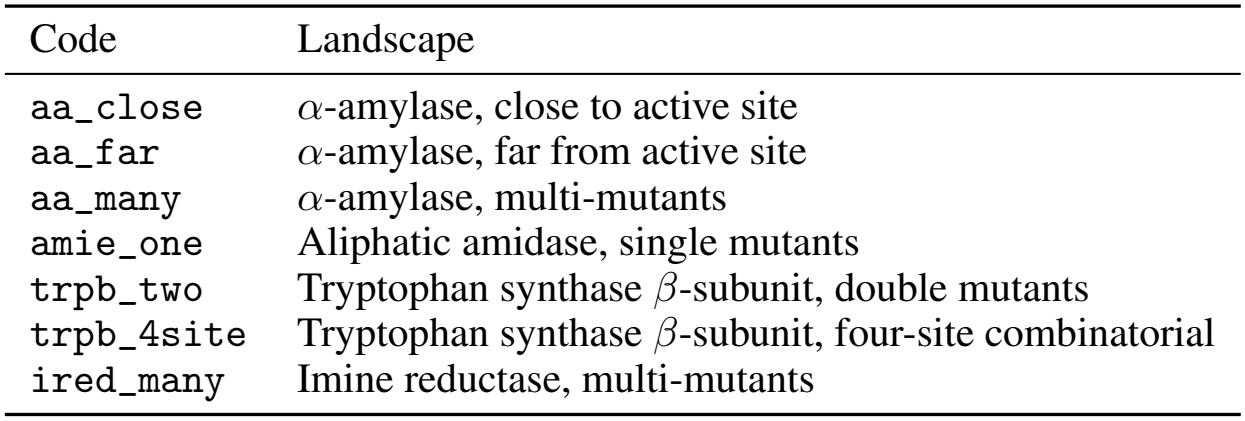
Mapping between the internal landscape codes used in Figure 6 and the corresponding landscape, matching the naming used in Figure 2 and Table 2.

**Table 4:** Discovery coverage of the top-5% of each landscape (†top-1% for TrpB 4-site) after 5 BO iterations using ESM2-650M as the PLM (96 acquisitions per batch), UCB (*β* = 1). Mean± SEM. over 10 seeds; best per row in bold.

| Landscape | One-hot | ESM2 / mean | finetuned ESM2 / mutpool |
|---|---|---|---|
| aa_close | 0.647 ± 0.011 | 0.506 ± 0.033 | **0.707 ± 0.031** |
| aa_far | 0.434 ± 0.007 | **0.653 ± 0.041** | 0.599 ± 0.016 |
| aa_many | 0.363 ± 0.021 | 0.229 ± 0.010 | **0.488 ± 0.028** |
| ired_many | 0.138 ± 0.009 | 0.151 ± 0.006 | **0.157 ± 0.007** |
| amie_one | 0.038 ± 0.006 | 0.127 ± 0.019 | **0.274 ± 0.008** |
| trpB_two | 0.377 ± 0.036 | 0.314 ± 0.008 | **0.431 ± 0.051** |
| trpB_4site<sup>†</sup> | 0.133 ± 0.025 | 0.030 ± 0.020 | **0.158 ± 0.010** |

**Table 5:** Top-5% coverage († top-1% for TrpB 4-site) after 5 rounds of BO, comparing static mean pooling, static mutation pooling, static context-aware pooling (*k*=4, *λ*=0.5, *τ* =1.0), and fine-tuned mutation pooling. Mean ± SEM over 10 seeds; best per landscape in **bold**.

| Landscape | static_esmc/mean | static_esmc/mut | static_esmc/context | finetuned_esmc/mut |
|---|---|---|---|---|
| aa_close | 0.471 ± 0.049 | 0.315 ± 0.013 | 0.477 ± 0.078 | **0.761 ± 0.019** |
| aa_far | 0.524 ± 0.028 | 0.264 ± 0.045 | 0.536 ± 0.026 | **0.705 ± 0.017** |
| aa_many | 0.299 ± 0.025 | 0.253 ± 0.028 | 0.340 ± 0.048 | **0.574 ± 0.015** |
| ired_many | 0.077 ± 0.002 | 0.000 ± 0.000 | 0.134 ± 0.003 | **0.169 ± 0.007** |
| amie_one | 0.093 ± 0.009 | 0.176 ± 0.006 | 0.220 ± 0.011 | **0.264 ± 0.006** |
| trpB_two | 0.122 ± 0.026 | 0.151 ± 0.017 | 0.229 ± 0.029 | **0.533 ± 0.016** |
| trpB_4site† | 0.001 ± 0.001 | 0.012 ± 0.007 | 0.029 ± 0.010 | **0.225 ± 0.016** |

**Figure 6.**
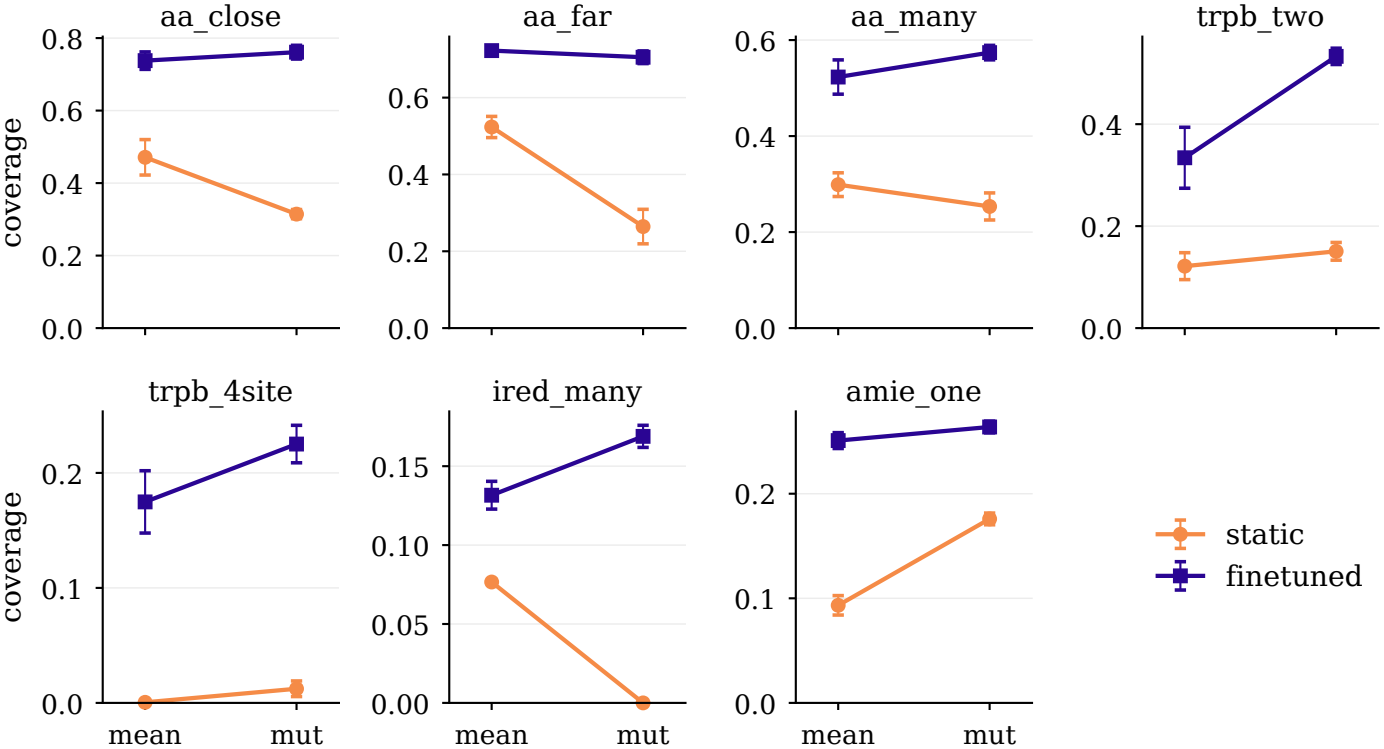
Top-5% coverage († top-1% for TrpB 4-site) after BO on seven benchmarks, for static vs. finetuned ESM-C under mean vs. mutation-site pooling. Points are means over 10 seeds, error bars SEM.; *y*-scale differs per panel.

## C Generalization

The generalization experiments use a train/test split of the alpha-amylase and tryptophan synthase *β*-subunit enzyme landscapes to test the out-of-distribution (OOD) capabilities of the trained surrogate (GP). We use four splits:

- Close→ Far: the GP is trained on sequences with mutations close to the active site (i.e., within 7.3Å of the active site), while the test set contains mutations in distant, unperturbed regions.
- Far →Close: the reverse of the above, trained on sequences with mutations far from the active site and tested on sequences with mutations close to it.
- One→ Many: trained on sequences with single mutations and tested on sequences with multiple (2-8) mutations distributed throughout the sequence.
- Two → Many: trained on sequences with 1 or 2 mutations and tested on multi-mutants.

We first run 5 iterations on the train split to acquire a total of 576 sequences from that distribution, then warm-start the GP with these acquired sequences and run acquisition over the test set. The cold-start variant instead trains the GP on 96 below-median-fitness sequences drawn from the test set itself, with acquisition likewise run on the test set. The transfer experiment below covers the three alpha-amylase splits; the TrpB split is reported in Table 1 only. Figure 3 highlights two key observations:

1. **Warm start: the finetuned representation dominates**. Given information from a different mutational context, the finetuned representation achieves better coverage than the baseline representations. It leads at every acquisition round across all three splits, with the clearest separation where the baselines plateau furthest below the top-5% band. Frozen ESM-C tracks one-hot closely in this regime, indicating that the gain stems from task-adaptive fine-tuning rather than from pre-trained embeddings alone.
2. **Finetuned arms benefit most from the warm start while only Phase-1 knowledge is available**. Comparing warm- and cold-start transfer shows a clear advantage to providing training data from a different mutational context, for all three representations and across all three splits. However, averaged over the three splits, this improvement over the cold-start schedule is largest at the first acquisition round, where the batch is selected by the Phase-1 model before any test-set label is observed; the baselines close part of the gap only once test-set feedback accumulates over later rounds. This advantage is therefore attributable to what fine-tuning transfers, rather than to the acquisition loop itself.

## D PLM Ablation

To test whether our conclusions are specific to ESM-C, we repeat the discovery experiments with ESM2-650M as the encoder, comparing one-hot encoding, static mean-pooled embeddings, and LoRA fine-tuning with mutation-site pooling (Table 4). The ranking of representation strategies is preserved: the finetuned arm attains the highest coverage on six of the seven landscapes, while one-hot encoding is best on none. The behavior observed with ESM-C on `amie_one` recurs here, with one-hot collapsing and fine-tuning recovering roughly sevenfold over it; on `trpB_4site`, static mean-pooled embeddings likewise fall below one-hot while the finetuned arm surpasses both. Absolute coverages are lower than those obtained with ESM-C on most landscapes, indicating that encoder quality continues to matter, but the advantage of adapting the representation is not tied to a particular PLM.

## E Mutation-aware pooling

Motivated by the strong performance of mutation-site pooling, we investigate whether incorporating local sequence context around mutated positions can further improve the representation. Pure mutation pooling discards all information from non-mutated residues, which may omit structurally or functionally coupled positions whose embeddings carry relevant signal. We therefore propose a context-aware extension that, for each mutated residue, identifies and incorporates the residues whose representations are most similar to it.

Concretely, for each mutated position *m* in a sequence, we compute the cosine similarity between the hidden representation at *m* and all other valid (non-mutated) positions. To encourage selection of spatially proximate residues, we apply a logarithmic sequence-distance penalty controlled by a locality parameter *λ*:

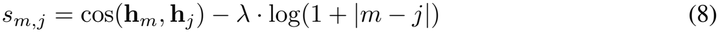

The top-*k* positions with the highest penalized similarity scores are selected, and a context vector for each mutation is obtained by softmax-pooling over the mutated residue itself and its *k* nearest context tokens:

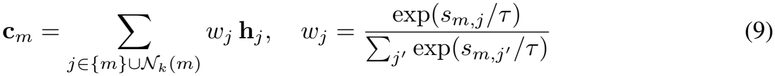

where *τ* is a temperature parameter and *N*_*k*_(*m*) denotes the top-*k* context set for mutation *m*. The per-mutation context vectors are then averaged across all mutated positions to produce a single fixed-dimensional sequence representation, as in standard mutation pooling. Setting *k*=0 recovers pure mutation-site pooling as a special case.

This formulation is fully differentiable with respect to the hidden states and is therefore compatible with LoRA-based fine-tuning: while the top-*k* selection itself is discrete, gradients flow through both the selected hidden states and their softmax weights into the encoder.

We find that incorporating mutational context improves the performance of static (frozen) baselines, narrowing the gap to fine-tuned representations. However, the static context-aware variant still falls short of the fine-tuned mutation pooling across all landscapes (Table 5), indicating that the primary source of improvement remains the adaptation of the representation itself rather than the pooling mechanism alone. We report these results primarily to document the direction; a systematic exploration of the interplay between context-aware pooling and joint fine-tuning is left to future work.

## F Pooling and representation ablation

Figure 6 shows that mean-pooling only the mutated residue positions collapses in the static (non-finetuned) GP setting, whereas in the LoRA regime, mutation pooling makes far better use of the adapted representation than mean-pooling the full sequence. This collapse in the static setting is expected: although the attention mechanism means each residue’s embedding reflects the influence of other residues, that influence encodes only evolutionary context. Mutated-site embeddings are therefore uninformative on their own, and fine-tuning is what transforms them into discriminative, function-relevant features. As the PLM comes to encode information specific to the target function being optimized, non-trivial pooling methods better isolate this mutational signal within the GP input. In the finetuned setting, even mean pooling provides a substantial advantage over the baselines, though mutation- and learned-pooling remain more consistent across benchmarks.

